# Assessing and Maximizing Cultivated Diversity with Plate-Wash PCR and High Throughput Sequencing

**DOI:** 10.1101/2020.11.19.390864

**Authors:** Emily N. Junkins, Bradley S. Stevenson

**Affiliations:** Department of Microbiology and Plant Biology, University of Oklahoma, Norman, OK, USA

**Author notes:** Address correspondence to Bradley S. Stevenson.

## Abstract

Molecular techniques continue to reveal a growing disparity between the immense diversity of microbial life and the small proportion that is in pure culture. The disparity, originally dubbed “the great plate count anomaly” by Staley and Konopka, has become even more vexing given our increased understanding of the importance of microbiomes to a host and the role of microorganisms in the vital biogeochemical functions of our biosphere. Searching for novel antimicrobial drug targets often focuses on screening a broad diversity of microorganisms. If diverse microorganisms are to be screened, they need to be cultivated. Recent innovative research has used molecular techniques to assess the efficacy of cultivation efforts, providing invaluable feedback to cultivation strategies for isolating targeted and/or novel microorganisms. Here, we aimed to determine the efficiency of cultivating representative microorganisms from a non-human, mammalian microbiome, identify those microorganisms, and determine the bioactivity of isolates. Molecular methods indicated that around 57% of the ASVs detected in the original inoculum were cultivated in our experiments, but nearly 53% of the total ASVs that were present in our cultivation experiments were *not* detected in the original inoculum. In light of our controls, our data suggests that when molecular tools were used to characterize our cultivation efforts, they provided a more complete, albeit more complex, understanding of which organisms were present compared to what was eventually cultivated. Lastly, about 3% of the isolates collected from our cultivation experiments showed inhibitory bioactivity against a multidrug-resistant pathogen panel, further highlighting the importance of informing and directing future cultivation efforts with molecular tools.

**Importance:** Cultivation is the definitive tool to understand a microorganism’s physiology, metabolism, and ecological role(s). Despite continuous efforts to hone this skill, researchers are still observing yet-to-be cultivated organisms through high-throughput sequencing studies. Here, we use the very same tool that highlights biodiversity to assess cultivation efficiency. When applied to drug discovery, where screening a vast number of isolates for bioactive metabolites is common, cultivating redundant organisms is a hindrance. However, we observed that cultivating in combination with molecular tools can expand the observed diversity of an environment and its community, potentially increasing the number of microorganisms to be screened for natural products.

## Introduction

Molecular techniques and sequencing technologies have helped to describe many microbial communities and provide tools that allow us to study representative microorganisms and their activities without isolation. Application of molecular approaches to the study of microbiology have led to the discovery and characterization of rare biospheres of the most extreme environments (1) and enabled researchers to decipher the metabolic potential and activity of microorganisms from the human gut (2) to the bottom of the ocean (3). Now, changes in microbial community structure and function can be monitored with molecular techniques throughout the course of a disease (4) or with changing environmental conditions (5), such that we may begin to predict the roles of microbial populations and outcomes over time (6, 7).

Isolating, or even just cultivating representative microorganisms is still considered the best means towards understanding their physiology, metabolism, and potential ecological roles, but this remains to be a major hurdle. The degree to which any microbial community is represented in culture varies considerably. Since Staley and Konopka coined the term “great plate count anomaly” (8), numerous studies have sought to quantify the culturable organisms from various biomes (see (9) for a compiled list of most probable number (MPN) based approaches). A recent study estimated that 81% of all microbial cells on Earth belong to uncultured genera or higher classifications and about 25% of microbial cells are from uncultured phyla (9). Specifically, 26% of marine, 31% of host-associated, 31% of eutrophic lakes, and 30% of soil metagenomic sequences have cultured representatives at the family level (9). Not surprisingly, human associated microbiomes have more cultured representatives, likely affected by a focused effort (10). Sequencing technology has shed light onto the previously unknown diversity of the microbial world, outpacing our ability to cultivate these organisms by an estimated rate of 2.4- and 2.5-fold for bacteria and archaea, respectively (11).

Efforts to isolate yet uncultured microorganisms have gathered momentum as researchers have begun to apply new technologies and innovations (12–14). Microbiologists routinely manipulate the composition of growth media and treatments of the inocula in order to increase the diversity of microorganisms represented in culture (e.g., (14–22)). As an example, treating an inoculum with ethanol can select for spore-forming bacteria by killing vegetative cells and leaving spores intact that would otherwise be overgrown and remain un-isolated (23). Furthermore, antioxidants in media can support the cultivation of both aerobic and anaerobic organisms, by using compounds that alleviate oxidative stress (24, 25). With the goal of this study being to cultivate many, different microorganisms, the approaches described above were of particular interest. We hypothesized that they would select for gut-associated bacteria; antioxidants would help mitigate oxygen stress for organisms best suited for microoxic or anoxic environments, and ethanol treatment would reduce the abundant vegetative cells, allowing us to better access the many spore-forming taxa that reside in the gut.

When cultivation approaches are coupled with molecular analyses, the total diversity and identity of cultivated taxa can be determined (26–28), and specific, targeted taxa can be detected (13). More recently, high throughput sequencing has made it possible to quantify exactly what proportion of the original sample has been cultivated (26, 29, 30). Studies using this approach showed that roughly 23% of microorganisms in bovine rumen fluid were cultivable in liquid enrichments (29), while 96% of those in human bronchoalveolar lavage samples (30), and 60% of those detected in/on American toads (26) were cultivated. A recent study characterized the microbial diversity within the same inoculum spread onto both high- and low-nutrient media by harvesting biomass on each plate and identifying all taxa present (26). Their approach built upon a previous study that sought to isolate abundant but yet-uncultured taxa using defined, low nutrient media and a PCR-based screening approach using group specific primers, termed platewash PCR (PWPCR) (13). Medina et al. confirmed that low nutrient-media would allow for the growth of a more diverse subset of bacteria from the original sample and that molecular screening (i.e., 16S PWPCR) is a practical way to characterize the microbial community growing on an entire agar plate (26). This data is invaluable in identifying medium composition, inoculum treatments, and incubation conditions that can target certain taxa, or maximize the cultivated diversity for large-scale isolation efforts (see 16, 18). However, these studies using solid agar plates may have overestimated diversity for some taxa by focusing on the relative abundance of taxa in their analyses (see Discussion).

Pure cultures have always been the “gold standard” for the in-depth study of microorganisms. The need for isolation is especially important when screening for the production of secondary metabolites for drug discovery. Historically, the process of isolating microorganisms for drug discovery has relied mainly on high throughput cultivation efforts that increase the number of isolates being screened (e.g., (31–33)), enrich for bioactive production from known producers like *Streptomycetes* spp. (34), or focus on novel or rare taxa thought to have uncharacterized biosynthetic capabilities (35–37). We aimed to do the same by sampling a diverse, host-associated microbial community and cultivating them under conditions designed to obtain as many different organisms as possible through the use of different media types. Screening cultivated microorganisms with the broadest diversity is expected to provide the greatest chance of uncovering natural products that are novel drug targets. Previous studies have focused on extreme environments for novel natural products (38), but we expand on this idea to include non-human mammals. We predicted that this would increase the diversity of microorganisms and compounds screened for useful antimicrobial compounds and argue that the shared life-history between a host and its host-associated microbial communities may produce compounds that are relatively less cytotoxic to the host. This idea was originally supported by a survey of microbiomes collected from roadkill mammals, which produced two new cyclic lipodepsipeptides of pharmaceutical interest (33).

Here, we sought to use both general and selective media along with various treatments to the inocula to collectively enrich for as many different microorganisms as possible from the oral and gut microbiomes of a roadkill mammal (racoon). The goals for this study were to (i) compare the diversity of cultured bacterial communities on multiple types of media combined with various treatments of the inoculum, (ii) determine which media, treatments, or combination cultivated the highest richness of bacterial taxa, and (iii) determine how many isolates produced bioactive molecules against a panel of multi-drug resistant pathogenic microorganisms.

## Materials and Methods

### Sample Collection and Storage

A 3 mile stretch of Oklahoma State Highway 9 in Norman, OK between the University of Oklahoma, Norman campus and Lake Thunderbird was used for our opportunistic sampling area. This well-traveled highway runs through undeveloped prairie, scrub and forest, cattle ranches, and rural residential land, providing many mammalian roadkill specimens. Sample collection was conducted under Oklahoma Scientific Collector Permit #5250 (33). The oral cavity and rectum of a racoon *(Procyon lotor)* deceased less than 8 h was sampled in triplicate (n=3) with sterile, nylon swabs. To ensure equal biomass was collected for immediate sequencing and subsequent cultivation, two swabs were used to sample each orifice simultaneously (to increase amount of sample) and stored together in a sterile conical tube containing 3.0 mL 1X PBS, stored on ice, and immediately transferred back to the laboratory (under 45 minutes). At the laboratory, each set of swabs was vortexed to suspend cells. A 900 μL aliquot of the cell suspension intended for molecular analysis was centrifuged at 10,000 x g for 1 min, the supernatant was removed. The cell pellet was resuspended in 750 mL of BashingBead Buffer (Zymo Research Corp.) and transferred to BashingBead Lysis Tubes (Zymo Research Corp., Irvine, CA) prior to lysis through homogenization in a Mini-BeadBeater-8 (BioSpec Products Inc., Bartlesville, OK) at maximum speed for 45 seconds. Homogenized samples were then stored at −20°C until needed for DNA extraction. For cultivation, the remaining 2.1 mL of the cell suspension for each sample was serially diluted with PBS. Aliquots of 50 μL from the 10^0^-10^-6^ dilutions were spread onto 13 different media/treatment combinations (see Table S1). This 2.1 mL volume represented the exact amount needed for cultivation with no suspension left over so that the 10^0^ dilution was not diluted and represented the surface area of the swab. In all, each inoculum was spread onto the agar media, in triplicate, for each media type and incubated for 6 days at 30°C.

### Culture-Dependent Bacterial Community Analysis

Cultivation was conducted with a combination of broad-spectrum media with decreased nutrients and selective media in order to “cast a wide net” and maximize the cultivable diversity of each inoculum (see quasi-factorial design in Table S1). The media used included 0.1X strength tryptic soy broth (Bacto, USA) with 1.5% agar, ROXY and derivatives with hemin (0.1 g/L) and alpha-ketoglutarate (2.0 g/L) (25), 0.25X strength R2A medium (15), and blood agar (per L: 10.0 g meat extract, 10.0 g peptone, 5.0 g sodium chloride, 15 g agar, 5% sheep’s blood (ThermoFisher, USA), pH 7). To date, this is the first instance using ROXY medium and plate-wash PCR techniques. In addition to different media types, one variation included the pretreatment of the inocula for 4 hours in 70% ethanol at 22-24°C (23). Catalase (40,000 U/L) was added as a treatment to TSA, ROXY and R2A to remove peroxides produced during autoclaving (19, 20). The last treatment included the addition of streptomycin at 50 mg/L as a broad spectrum selection agent against Gram-negative and some Gram-positive species (39).

To test which combinations of media and treatments resulted in cultivated communities that represented the diversity observed in the original sample, we grew each inoculum (mouth or rectum) on each combination of media/treatment types and sequenced the resulting biomass.

### Collection of Total Biomass from Agar Plates, DNA Extraction and PCR (PWPCR)

Biomass was harvested from each agar plate in order to identify the microorganisms present and to compare this to culture-independent analyses of the inoculum. The biomass from each agar plate was harvested by adding 2.0 mL of PBS to the surface of the plate and suspending the colony biomass by agitation with a sterile spreader. The suspended colony biomass was collected and transferred to a 2.0 mL microcentrifuge tube, pelleted at 10,000 x g for 1 min, and resuspended in 750 uL of BashingBead Buffer (Zymo Research Corp.). Each sample was transferred to a BashingBead Lysis Tube (Zymo Research Corp.) and homogenized for 45 seconds at maximum speed using a Mini-BeadBeater-8 (BioSpec Products Inc., Bartlesville, OK). The lysed samples were stored at −20°C until DNA was extracted.

### DNA Extraction and Sequencing

Before DNA extraction, each sample was thawed on ice and homogenized for 30 seconds at maximum speed using a Mini-BeadBeater-8 (BioSpec Products Inc., Bartlesville, OK). DNA was extracted according to manufacturer specifications using the Zymo *Quick-DNA* Fungal/Bacterial Miniprep kit (Cat# D6005, Zymo Research Corp., Irvine, CA). For community characterization, a conserved region of the SSU rRNA gene of most bacteria, archaea, and eukarya was amplified using primers 515F-Y and 926R (40) via the following PCR protocol: initial denaturation at 94°C for 2 min, followed by 30 cycles of denaturation at 94°C for 45 s, annealing at 50°C for 45 s, and extension at 68°C for 90 s, with a final extension at 68°C for 5 min. These primers produced two amplicons, a ~400 bp fragment for bacteria and archaea, and a 600 bp fragment for eukaryotes. The forward primer 515F-Y (5’-<u>GTA AAA CGA CGG CCA G</u> CCG TGY CAG CMG CCG CGG TAA-3’) contains the M13 tag (underlined) fused to the 5’ end of the forward primer (41). The reverse primer 926R (5’-CCG YCA ATT YMT TTR AGT TT-3’) was unmodified from Parada et. al 2015. Each PCR contained 5 PRIME HOT master mix (1X; 5 PRIME Inc., Gaithersburg, MD), 0.2 μM of each primer, and 3.0 μL of extracted DNA at a final volume of 50 μL. The amplified fragments were purified using Sera-Mag magnetic beads (GE) with the AmPureXP (Beckman Coulter) protocol at a final concentration of 1.8x v/v. After purification, 3 μL of each PCR product, 1x 5 PRIME HOT master mix (Quantabio, Massachusetts, USA), 0.2μM of the 926R primer, and 0.2 μM of a specific 12 bp oligonucleotide was used in a separate barcoding PCR (6 cycles) in 50 μL reactions to attach a unique barcode to amplicons of each library. The same thermocycler protocol was used as above but only run for 6 cycles. The now barcoded amplicons were purified using Sera-Mag (GE) beads with the AmPureXP (Beckman Coulter) protocol to a final volume of 40 μL, quantified using the QuBit HS DS DNA assay kit (Thermo Fisher Scientific Inc., Waltham, MA), and pooled in equimolar amounts. The pooled, barcoded amplicon libraries were then concentrated to a final volume of 40 μL (209 ng/ μL) with an Amicon-Ultra filter (Millipore, Burlington, MA, USA) following manufacturer’s protocol. The combined amplicon libraries were denatured according to Miseq library preparations protocol (Illumina, San Diego, CA, USA). The sample was loaded at a concentration of 10 pM and sequenced using 2×250 paired-end strategy on the Miseq (Illumina San Diego, CA, USA) platform for 251 cycles.

### ASV Inference and Bacterial Community Characterization

Barcodes from raw SSU rRNA amplicon sequences were removed and demultiplexed using QIIME v 1.9.1 (42). Demultiplexed reads were trimmed for adapters and quality filtered as a part of the *dada2* pipeline (43) and amplicon sequence variants (ASVs) were inferred using the forward reads (203 bp). Taxonomy was assigned using the SILVA database v32 (44, 45). The final dataset consisted of ~1.7 million reads resulting in 463 ASVs with a median of 17,347 sequences per sample (min = 132, max= 266,678).

#### Data Availability

Sequence data has been deposited at NCBI’s Sequence Read Archive (SRA) database under accession number <u>PRJNA675861</u>.

### Data Analysis

Community diversity was analyzed using the *phyloseq* (46) and *vegan* (47) packages in R. Relative abundance was only used to directly assess the diversity of the original inocula. Presence/absence and richness were used as the main metrics to compare cultivated samples because of the variation in biomass (i.e. colony size) between microbial colonies on and between agar plates. Also, because of the sampling procedure with duplicate swabs used in triplicate, the data from the molecular control was pooled for analysis since the variability in swabbing would be different than the variability of triplicates resulting from cultivation.

Alpha diversity was calculated using richness of ASVs, while beta diversity was measured with non-metric multidimentional scaling (NMDS) using Jaccard distances. Statistical comparisons across media types, treatment, and orifices where done using permanova with the *adonis2* function in the *vegan* package (47). Due to the nestedness of orifice and cultivation conditions, orifice was used as a strata or fixed factor, and media and treatment were used as predictor variables (~media*treatment). Communities were also compared without using orifice as a fixed factor to capture any influence that orifice could have on the cultivable communities. To fully understand the significance described in the permanova, a permdisp with the *betadisper* function in *vegan* was used to describe any within sample variance (i.e. across replicate agar plates) that could explain any significant differences detected in the permanova. Hypothesis testing via anova with permutations (n=999) was used with permdisp to determine any significant differences in variation within samples.

To identify certain ASV families more associated with an orifice, medium, or treatment, a species indicator analysis was performed using the *indicspecies* package (permutations = 999) (48, 49).

### Bioassays and Isolate Identification

Colonies were selected for isolation based on morphology, with the intent of sampling as many different colony morphologies as possible. From average of 8.1 x 10^10^ CFUs/mL cultivated across all medium types and dilutions, a total 238 colonies were chosen for isolation based on differential colony morphology from the mouth and rectum on a subset of the media types (0.1X TSA, ROXY with alpha-ketoglutarate and hemin, and 0.25X R2A). To determine bioactivity, soft agar overlays in 0.1X TSA (0.8% agar) were used to test for growth inhibition of a panel of drug-resistant pathogens *(Klebsiella pneumonia* ATCC# 13883, *Enterococcus faecium* ATCC# 51559, *Pseudomonas aeruginosa* ATCC# 10145, and *Candida albicans* ATCC# 565304). To prepare for the bioassay, pathogens were incubated overnight, shaking at 250 rpm at 30°C, in 0.1X tryptic soy broth (TSB) (Bacto, USA). A 400 μL aliquot of each pathogen was used to inoculate 3.6 mL of molten TSA soft agar (at 55 °C), mixed, and immediately poured onto a 0.1X TSA agar plate for a final overlay volume of 4 mL. Colony material from each isolate was transferred to the bioassay plate with a sterile toothpick by etching an ‘X’ into the overlay. The inoculated bioassay plates were incubated at 30 °C for 24 hours. Inhibition of the pathogen in the overlay was characterized by a zone of clearing surrounding the isolate (Figure S1). For cryopreservation, each isolate was grown in 5 mL of 0.1X TSB in a 16 x 150 mm test tube shaking at 250 rpm for 48 h. An aliquot of 800 uL of the culture was transferred to a 2 mL screw cap tube, along with 200 uL of 80% glycerol, for a final concentration of 20% glycerol. After mixing by vortex for 30 s, it was then stored at −80°C.

Partial SSU rRNA sequences were used to identify each isolate. Genomic DNA was extracted from each isolate using QuickExtract (Lucigen, Wisconsin, USA) according to manufacturer’s protocol from the above mentioned 0.1X TSB cultures. The SSU rRNA gene was amplified using primers 8F (3’-ATGC-5’) and 1492R (3’-ATGC-5’) at a concentration of 0.2 μM, 1x 5 PRIME HOT master mix (Quantabio, Massachusetts, USA), and 2.0 μL of DNA template per 50 μL reaction via the following PCR protocol: initial denaturation at 94°C for 2 min, followed by 30 cycles of denaturation at 94°C for 30 s, annealing at 55°C for 45 s, and extension at 72°C for 45 s, with a final extension at 72°C for 10 min. Amplified fragments were purified using Sera-Mag magnetic beads (GE) with the AmPureXP (Beckman Coulter) protocol at a final concentration of 1.8x v/v. Purified amplicons (a total of 192 samples) were sent for Sanger sequencing using the 8F primer (Genewiz, New Jersey, USA). Resulting sequences and chromatograms were assessed for quality using IGV (50). Sequences with low quality (< Q20) were removed, low quality ends were trimmed with MEGA X (51), and the final sequences were identified using the SILVA v. 132 classifier online server (44). A total of 117 isolates had suitable reads and met quality thresholds for taxonomic identification.

## Results

### Cultivated Microbial Diversity Differed Between Orifice and Cultivation Condition

As expected, a library of SSU rRNA genes from the original inocula showed that it had higher richness (alpha diversity) than the cultivated organisms from both orifices (Figure 1). The microbial community sampled from the rectum had higher overall richness (153 ASVs) compared to that of the mouth (86 ASVs) (Figure 1). Overall, we were able to cultivate 57.3% of the ASVs detected in the inocula (58.1% from the mouth and 57.5% from the rectum).

**Figure 1:**
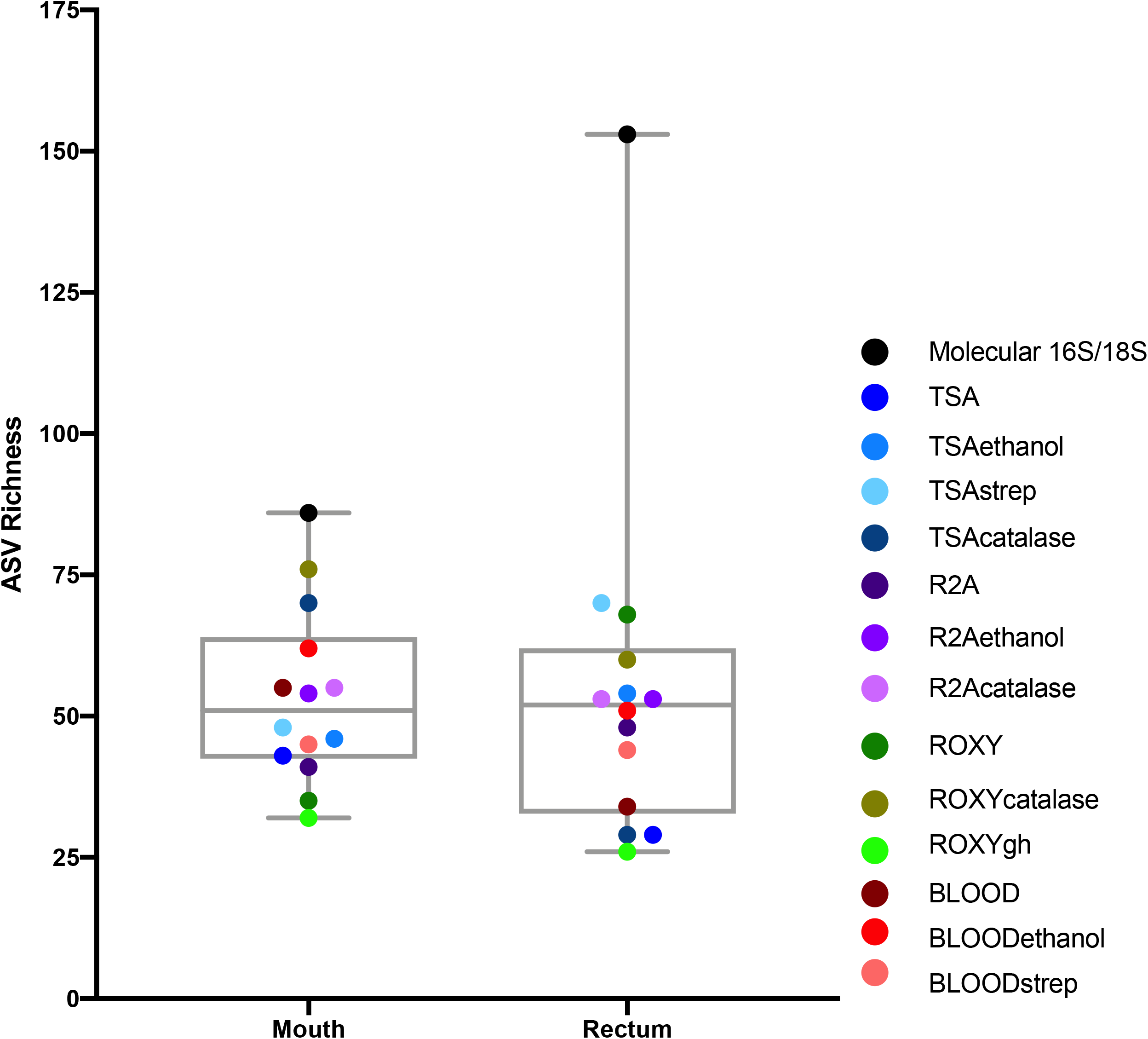
Alpha diversity of the mouth and rectum. For the controls, richness, measured using ASVs, was higher in the rectum than the mouth. For cultivated communities, richness varied based on medium type and treatment. Points represent each medium or treatment type, indicated by color (n=3). Molecular 16S/18S refers to the directly sequenced inoculum sample (n=3).

Interestingly, about half (52.7%) of the ASVs detected from the cultivation experiments were not represented in either library from the inocula (i.e. molecular control) (Figure 2). The number of ASVs detected on a certain medium or treatment differed between orifices. For instance, cultivation on TSA medium treated with catalase resulted in only 29 ASVs from the rectum inoculum compared to around 70 different ASVs from the mouth. The fewest number of ASVs was detected on the ROXY medium amended with alpha-ketoglutarate and hemin from both the mouth and rectum. The cultivated beta diversity differed based on orifice and cultivation condition. Specifically, community structure differed based on orifice and treatments, more so than base medium type (Figure 3). Differences in the cultivated community composition and diversity between medium and treatment confirmed that, collectively, using different cultivation techniques increased the number of organisms cultivated from each inoculum.

**Figure 2:**
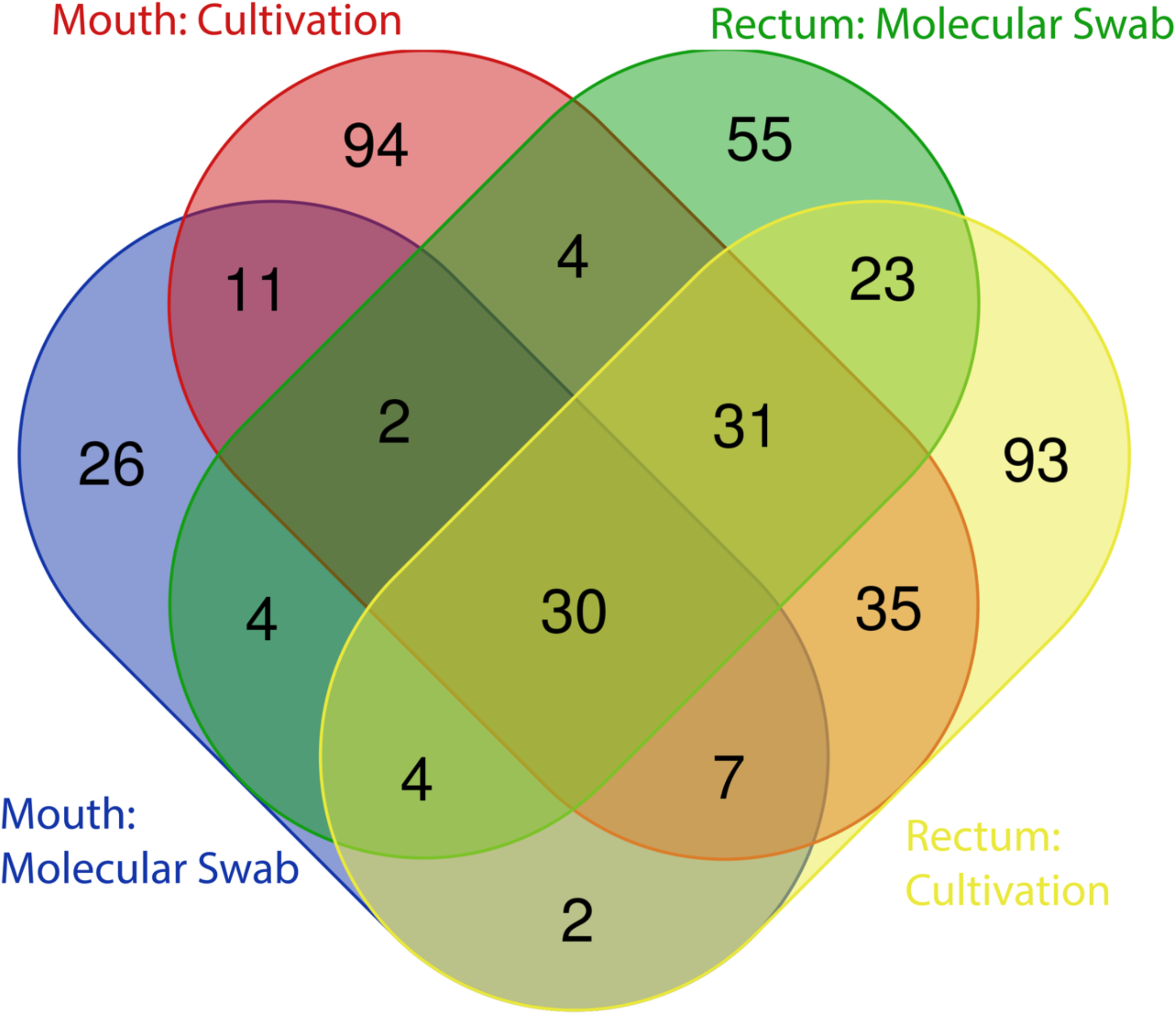
Distribution of ASVs between cultivation and molecular controls. The majority of ASVs detected in the original inocula were shared between the mouth and rectum, while approximately half of all ASVs detected through cultivation were not detected in the directly sequenced samples.

**Figure 3:**
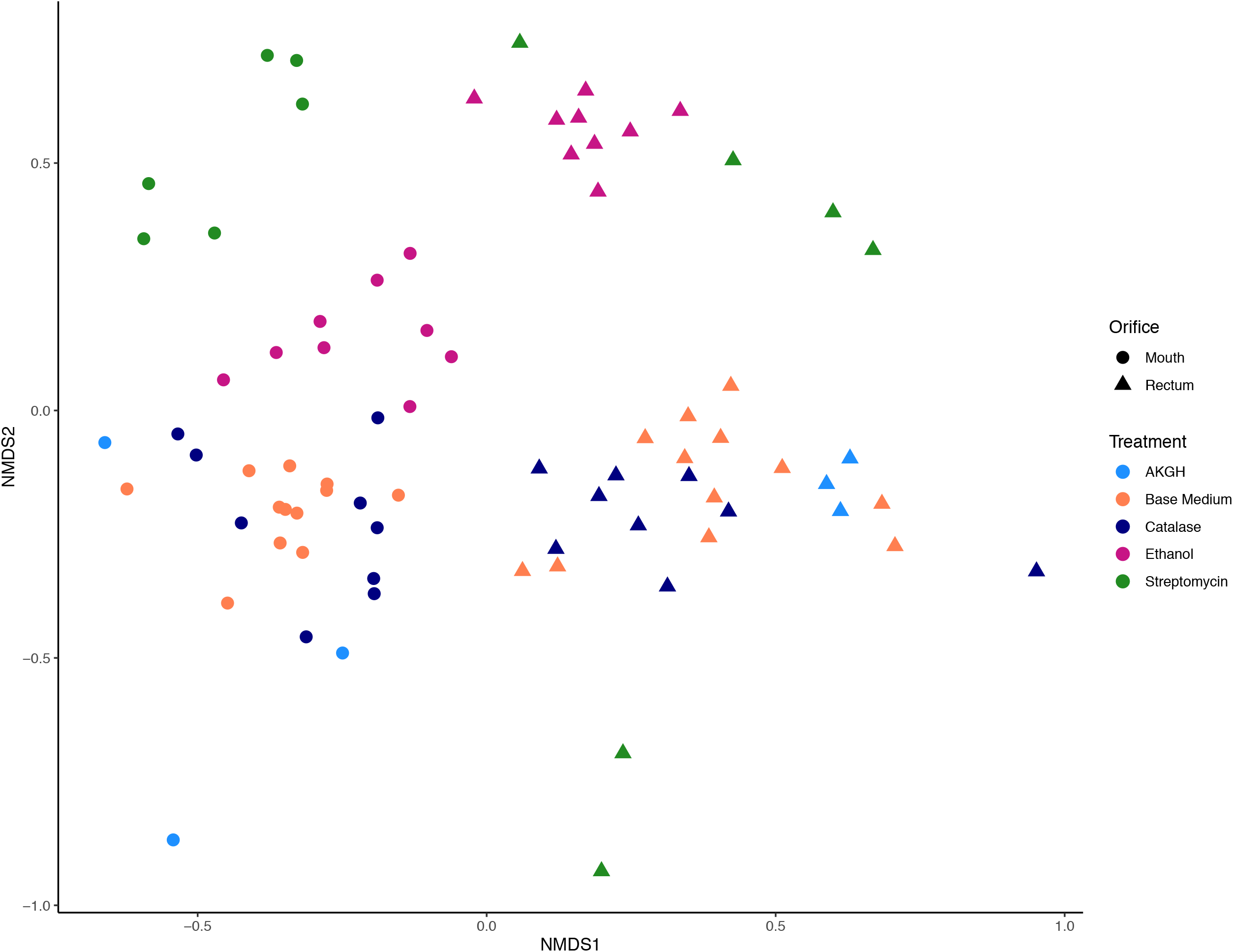
Beta diversity measured in Jaccard distance and NMDS ordination. Differences in microbial community structure were driven by orifice and treatment, specifically ethanol and streptomycin. Shape corresponds to orifice sampled, while color differentiates between treatments of the medium or inoculum.

### Selective Treatments Increased Cultivated Richness Compared to Media Type or Orifice

An indicator species analysis was used to determine which microbial families were significantly associated with certain conditions. A total of ten families were identified to be significantly (p < 0.05) associated with orifice, media type, and/or treatment. Only one family, Aeromonadaceae was significantly associated with a particular orifice, the mouth. This family was also the only family significantly (p = 0.001) associated with a certain base medium type, ROXY (Table 1). Treatment of the medium and/or the inoculum selected for the most diverse set of indicator species, with a total of 9 families significantly (p < 0.05) associated with a certain treatment or treatments (Table 1). Certain bacterial families like Staphylococcaceae and Paenibacillaceae were associated with the ethanol pretreatment of inocula, whereas the Wohlfarhtimonodaceae and Metaschinikowiceae were more associated with the addition of streptomycin to media (Table 1).

**Table 1.**
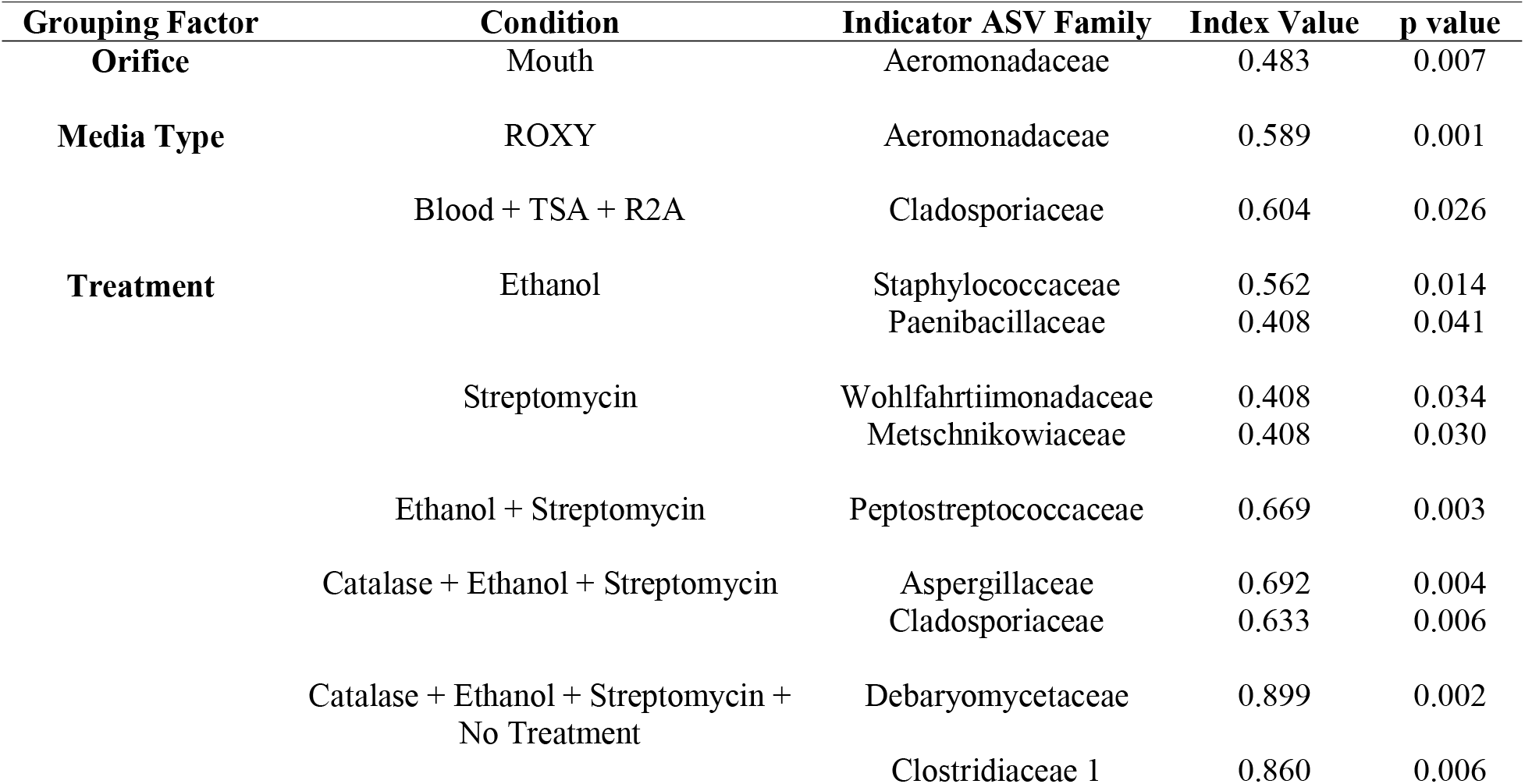
Taxonomic associations with media type using Indicator Species Analysis

Despite some unique taxa being associated with certain conditions, much of the cultivable microbial diversity was shared between the mouth and the rectum (Figure 4), indicated by the lack of significant differences in community structure described by the permanova and permdisp analyses (discussed below). Based on molecular analyses, 27 families where shared between the mouth and rectum before cultivation, while 6 families were unique to the mouth and 6 families were unique to the rectum. Negative cultivation controls (e.g., PBS) yielded no growth, supporting the assumption that any biomass collected from plates originated from the inoculum. Lastly, our cultivation experiments led to the detection of 18 families from the mouth and 8 families from the rectum that were not detected with direct molecular analysis of the inoculum. This accounted for around half (52.7%) of the ASVs detected during cultivation (Figure 2). This could be explained by differences in biomass and growth characteristics not being captured by sequence data from the inocula (discussed below).

**Figure 4:**
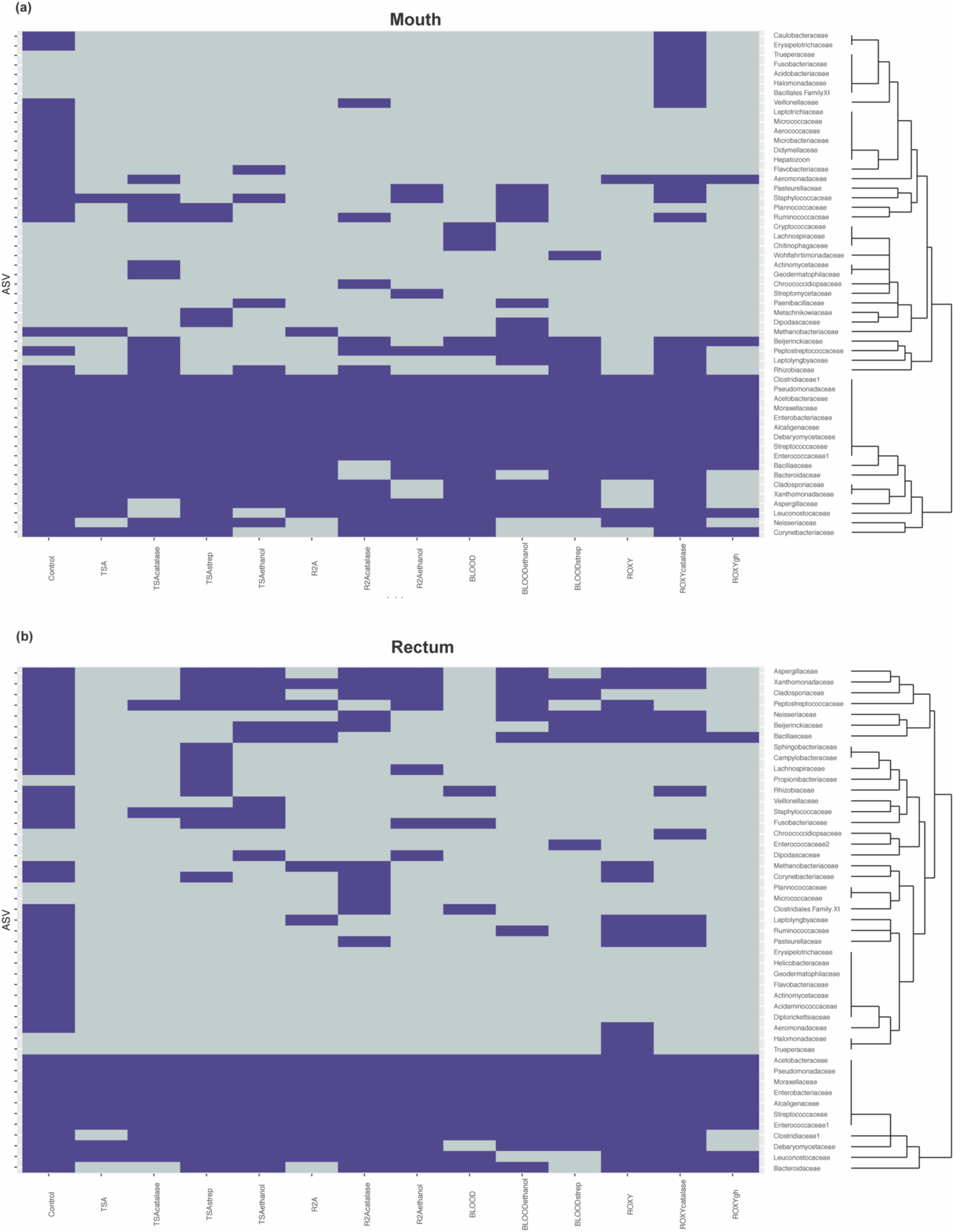
Heatmaps describing presence absence of taxonomic families observed in the (a) mouth and (b) rectum. Purple indicates that a family was observed while grey indicates that a family was absent. The dendrograms highlight clusters of families observed during cultivation.

### The Raccoon Microbiome Contained Cultivable Bioactive Bacteria

A total of 238 isolates were collected from a subset of media (0.25X R2A, 0.1X TSA, and ROXY with alpha-ketoglutarate and hemin). Many of these isolates were expected to be redundant, since the diversity of the initial inoculum was low (Figure 1). This was confirmed with many isolates redundant at the genus level and the majority of the identifiable isolates belonging to genera *Serratia* (34.2%) and *Klebsiella* (17.9%) (Figure S2). Each of the 238 isolates was assayed for antimicrobial activity against a panel of multi-drug resistant pathogens. A total of 7 isolates showed antimicrobial activity, identified as a zone of clearing of one of the antibiotic resistant panel organisms. This equated to a ~3% “hit” rate against already resistant organisms. Six of these isolates were recovered from the mouth, while one, a *Bacillus* sp., was recovered from the rectum. Three isolates were recovered from R2A, 3 from TSA and 1, an Enterobacteriaceae sp., from ROXYakgh. Three showed activity against *Klebsiella pneumonia,* three against *Enterococcus faecium,* and one, a *Pseudomonas* sp., against *Candida albicans.* Five of the isolates were identified by partial 16S rRNA sequence identity (Table 2).

**Table 2.** Cultivated bioactive isolates

| Isolate | SILVA Taxonomy | Orifice | Medium | Activity Against |
| --- | --- | --- | --- | --- |
| RC2RCR2A30-212 | *Bacillus* sp. | Rectum | R2A | *Klebsiella pneumoniae* |
| RC2MOR2A30-238 | *Pseudomonas* sp. | Mouth | R2A | *Candida albicans* |
| RC2MORGH30-018 | Enterobacteriaceae | Mouth | ROXYgh | *Enterococcus faecium* |
| RC2MOTSA30-050 | *Enterococcus* sp. | Mouth | TSA | *Enterococcus faecium* |
| RC2MOTSA30-142 | *Klebsiella* sp. | Mouth | TSA | *Klebsiella pneumoniae* |

### The Composition of Cultivated Taxa was Highly Variable

Due to the nestedness of this experiment (i.e. taxa observed during cultivation theoretically would be observed in sequencing), the stratification of taxa based on sampled orifice was corrected for hypothesis testing with permanova by defining orifice as a stratum in the *adonis2* function in the *vegan* package (47). The majority of the explained variation and significant differences in cultivated taxa were due to treatment of the inoculm or medium type (F= 2.44, R^2^ = 0.12, p = 0.001), while base medium type alone explained only ~5% of the observed variation (F= 1.54, R^2^ = 0.05, p = 0.002)(Figure 5). The interactions of base medium and treatment only explained ~6% of the variation (F= 1.52, R^2^= 0.06, p= 0.208). An analysis of variances (permdisp) was used to determine if any significance was driven by a dispersion effect rather than a location effect for orifice, base medium type, treatments, and medium+treatments. Mean distances to centroids did not differ significantly for orifice (F=0.69, p=0.41; Mouth 0.51; Rectum 0.53) or medium+treatment (F= 1.38, p=0.20; TSA=0.47, TSAcatalase=0.52, TSAstrep=0.51, TSAethanol=0.45, R2A=0.44, R2Acatalase=0.47, R2Aethanol=0.45, Blood=0.45, Bloodethanol=0.50, Bloodstrep=0.53, ROXY=0.48, ROXYcatalase=0.49, ROXYgh=0.52), meaning little inter-sample variation was detected between replicates. In contrast, the mean distances did differ for treatment (F=3.3, p=0.02; alpha-ketoglutarate+hemin=0.52, no treatment (base)=0.50, catalase=0.53, ethanol=0.49, streptomycin=0.56) and base medium type (F=4.4, p=0.01; Blood=0.56, R2A=0.50, ROXY=0.53, TSA=0.55). The significance in dispersion within the treatment and media could correspond to the significance in permanova results, where much of the variation explained in the permanova may be due to the intrinsic variability and stochasticity among replicates of the same inoculum.

**Figure 5:**
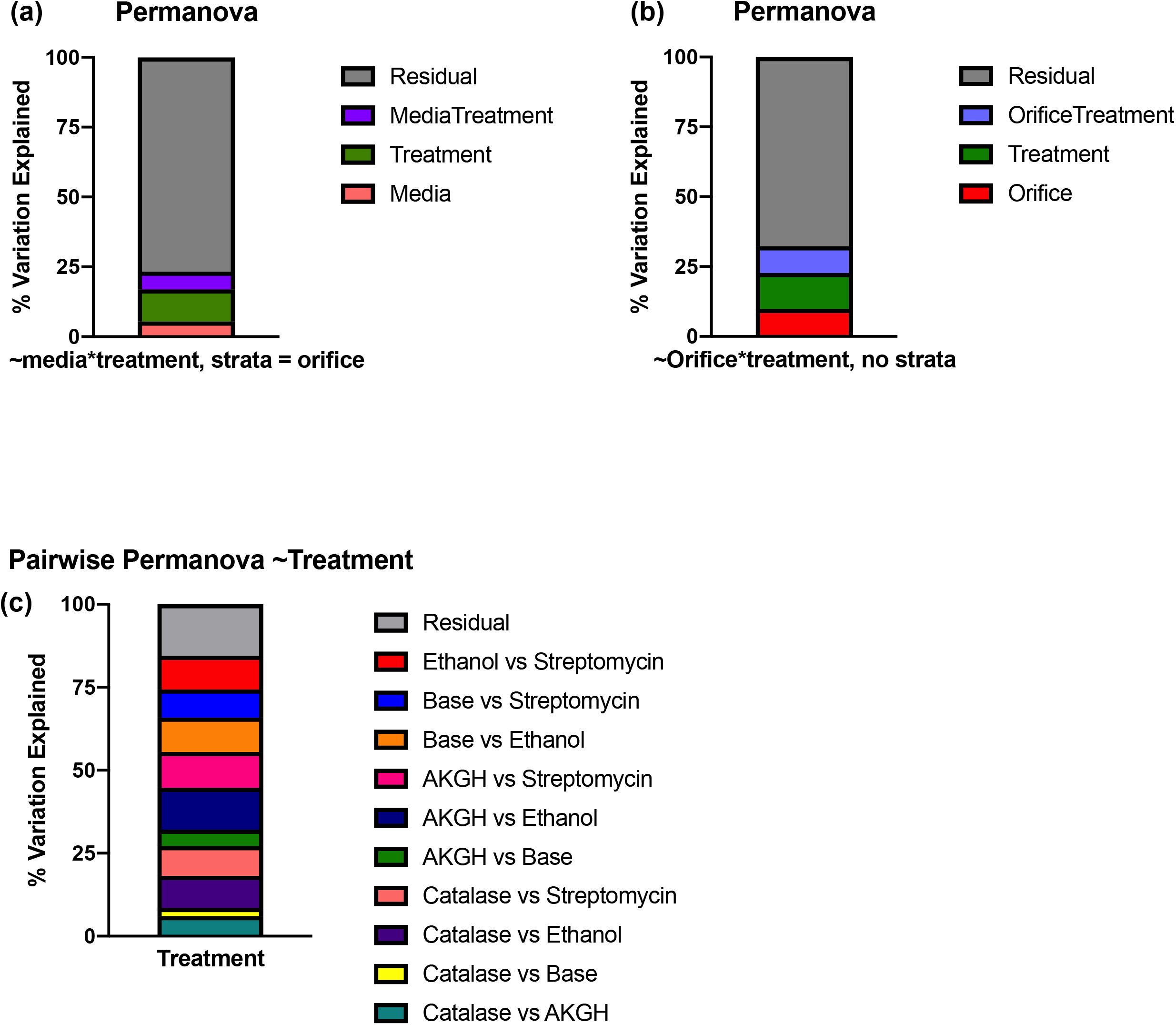
Percent of variation described by permanova models based on (a) media and treatment with orifice as a stratum, (b) orifice and treatment, and (c) pairwise comparisons of treatments on media types. Largely, the differences in community structure during cultivation could not be explained by media, orifice, or treatment alone. In all graphs, residual variation (in grey), is the proportion of variance not explained by the model.

### Discussion

We used mouth and rectal samples from a roadkill mammal (raccoon, *Procyon lotor)* to (1) compare the diversity of cultured bacterial communities on multiple media types and/or treatments of the inoculum, (2) determine which media/treatments cultivated the highest richness of bacterial taxa, and (3) determine if any isolated organisms produced bioactive molecules against a drug resistant pathogen panel. The occurrence of antimicrobial resistance is increasing at a faster rate than new therapies are entering the market (52, 53). Since nearly two-thirds of our current antimicrobial drugs are linked to microorganisms (52, 54), one of our goals was to cultivate a diverse array of bacteria from a non-human mammalian microbiome and screen these isolates for the production of antimicrobial metabolites. Growing microbial populations in the laboratory that are representative of the overall microbial diversity within an inoculum continues to be a major limitation for the field of microbiology. Here, we sought to use a combination of selective medium types and treatments of the inocula that might enrich for certain groups present in the oral and gut microbiomes of a roadkill racoon. In order to capture a more comprehensive sample of microbial diversity, we used high-throughput sequencing of 16S rRNA gene amplified libraries to characterize the original inocula and the cultivated populations that were washed from the surface of agar media.

We anticipated finding a significantly lower percentage of cultivable organisms relative to the molecular-based measures of diversity from each orifice, which we described as the cultivated percentage. We also anticipated that the identity of recovered taxa would be similar between media types and even orifices. The differences between organisms cultivated on different media, from different orifices was low. However, when observing the significance described in the permanova and permdisp, the percentage of explained variation was low. This suggested that the stoichiometry of diverse and variable inocula could play a large part in the outcome of this and any similar cultivation experiment. Based on the low percentage of explained variation, we also approached this data set from the perspective of the unobserved or residual variation, such that some of this discussion will aim to pose questions about what makes cultivation so difficult. Overall, we observed that cultivation yielded not just a fraction of what is detected in sequence data, but rather expanded the total diversity observed in the original sample.

### Cultivation can be optimized at large scales

Convergence of microbial communities occurred between the mouth and rectum for both the molecular samples and the cultivated communities. This was expected since as an animal undergoes decomposition, the microbial diversity decreases, especially in rectal communities (55), and becomes dominated by only certain taxa, namely Proteobacteria (56). In our samples, despite being a single time point roughly 8 hours after death, the mouth was dominated by Proteobacteria while the rectum samples contained mostly Firmicutes, Proteobacteria, and Bacteroides (Figure S3). Overall, about 87% of the microbial families detected in the molecular controls were shared between the mouth and rectum. This increase in shared taxa was also expected, as the proportion of cultivable organisms was expected to be lower (57) and the differences in diversity would not be detected through the selection of cultivability. Because of the small proportion of molecularly characterized microorganisms that are routinely recovered in the laboratory, we employed multiple variations in cultivation, like medium type and inoculum treatments to “cast a wider net.”

Treatments selected for organisms that can withstand ethanol stress (e.g., spore-formers), or were streptomycin resistant. These had a greater effect on the cultivable diversity compared to changes in base medium alone, likely because the base media shared many of the same oxidizable substrates (see media recipes in methods section). It was logical to hypothesize that the inoculum exposed to selective treatments could have decreased cultivated richness compared to the same inoculum on an untreated medium of the same type, since the selective agent would inhibit the growth of a portion of the population. Instead, the cultivable richness and diversity varied for the same treatments on different base medium types and between orifices (Figure 1 and Figure 5). This could be due to the treatments selecting for specific groups of taxa (i.e. antibiotics) or creating an environment where certain taxa were able to out-compete on non-selective medium (i.e. low nutrients, increased incubation and slow growers) (17). Specifically, medium amended with catalase or streptomycin and inocula treated with ethanol resulted in cultivated organisms not identified on other plates, or even the molecular controls in some cases (Figure 4). One possible explanation would be that this was due to the inherit stochasticity associated with cultivation (discussed below). Despite the variability, we were able to cultivate roughly 57% of the species richness observed with molecular approaches. Previous studies measuring cultivation success with high-throughput sequencing have reported similar levels of cultivable richness from the skin of toads (~60%) (26) and rumen fluid (23%) (29). Medina et al. (2017) reported greater variation in cultivability between inocula (different animals). Our data showed that more variability in cultivated taxa was observed between selective additions to the medium or treatments of the inocula (Figure 5), and not the inocula (different orifices) (Figure 3). Similar to Zehavi et al, (2018), different medium types and treatments increased the cultivable proportion of the original sample.

### Bioactivity and phylogenetic diversity of isolates

The research reported here was based on previous success of using roadkill mammals as a source of antimicrobial-producing microorganisms (33). Our intent was to quantify and broaden the diversity of cultivated bacteria by the addition of various media and treatments. This affect was measured by using plate-wash PCR and SSU rRNA gene sequencing. Simultaneously, colonies were also picked for isolation from replicate plates (n = 3), identified through SSU rRNA sequencing, and screened for antimicrobial production. Overall, 7 of the 238 recovered isolates showed bioactivity against the antibiotic-resistant pathogen panel, resulting in a ~3% hit rate. A total of five bioactive isolates were successfully sequenced and met our quality control standards described in the methods. Three belonged to the phylum Proteobacteria (class Gammaproteobacteria) and two belonged to the phylum Firmicutes. Two isolates were identified as a *Bacillus* sp. and a *Pseudomonas* sp., both genera known to contain species that produce bioactive compounds (33, 58–61). Interestingly, the isolates identified as an *Enterococcus* sp. and *Klebsiella* sp. showed inhibitory activity to a member of its own genus, *Enterococcus faecium* and *Klebsiella pneumonia,* respectively. This finding was consistent with the hypothesis that closely-related organisms often have a negative effect on each other due to resource overlap (62), which has been demonstrated for these genera (63). In addition, some populations could be making antimicrobials, like bacteriocins, that effect close relatives (64).

### Cultivation Yields Complex Data

The cultivation and isolation of many organisms at once can be incredibly laborious. New technologies and approaches can help increase throughput, which is necessary to overcome the re-isolation of abundant, common, and easy to cultivate organisms. However, it is still important to be able to assess the efficacy of any cultivation effort. Unfortunately, we uncovered an intrinsic property of cultivation that is not readily realized unless assessed at a large scale, variability. Here, we did not detect the same cells (e.g., colonies measured in PWPCR) consistently growing on replicate plates (Figure S4).

Neither differences in cultivation strategy nor orifice sampled could explain the majority of the observed variation in community structure (Figure 5). We can attribute unexplained variance (i.e. residuals) observed between communities (i.e. beta diversity) to stochastic effects of cultivation, but that is not fundamentally measurable in this case. We can speculate that randomness would play a role in cultivation as samples are diluted (27, 29) or the spatial heterogeneity of the inoculum is altered, as demonstrated on a much larger scale using leaf litter (65). In these instances, we could expect different inocula resulting in different assemblages of cultivated organisms. We also know that microorganisms immobilized on an agar surface are still able to interact through motility or diffusion of metabolites, making certain competitive adaptations more advantageous (64). In this experiment, the agar media allowed physical separation of individual cells in the inocula and constrained microbial populations to form colonies, potentially forcing interactions with neighboring colonies that could result in competition and inhibition of growth through secondary metabolite production (64). The randomness of which populations are in close proximity to one another could play a role in which members will thrive from the same inoculum source, affecting the outcome of which taxa are cultivated (66).

Cultivation on agar plates warrants special consideration when discussing diversity. The relative distribution of populations on an agar surface affects their prevalence and the overall composition of the cultivated community. Further, the physiology and colony morphology of each population dictates their prevalence and that of the surrounding populations on an agar surface. Colony size, especially surface area, is impacted by traits such as growth rate, surface motility (i.e., gliding motility, swarming), presence of inhibitory metabolites, and the availability of resources. These differences in biomass distribution were important when deciding how to measure diversity, a critical metric for assessing cultivation on an agar surface. Unlike liquid cultures, varying biomass (i.e. colony size) from different taxa that might produce an equal number of colonies can over-estimate the relative abundance of that taxon, studies relying on relative abundance to characterize diversity should consider this (26, 30). If a population (A) has more proficient growth and forms a larger colony than population (B), DNA extracted from each population would indicate that taxon A was more abundant in the original inoculum, incorrectly representing the evenness of the original inoculum. This is especially true when relative abundance is calculated based on equimolar DNA concentration during library prep. To mitigate any biased evenness, species richness and Jaccard distances was used to measure alpha and beta diversity, respectively.

In this data set, about half of the ASVs detected after cultivation were not detected in the molecular controls. This same phenomenon was observed by Zehavi et al. 2018 when cultivated rumen OTUs outnumbered the OTUs detected in the original rumen sample and its dilutions (1,012 out of 1,698) (29). This discrepancy may speak to the power of cultivation relative to the power of direct molecular analyses in describing the diversity of a community. However, both of these approaches come with their own caveats. Cultivation makes assumptions about an organism’s ability to grow in the laboratory, while sequencing is dependent on methodology, sequencing chemistry and sampling depth. In this study, rarefaction curves indicated that sequencing efforts were sufficient (Figure S5). One potential explanation for the large proportion of ASVs not detected in the controls could be that in combination with the different media and treatments, we were able to give some of the less abundant microorganisms a growth advantage. For example, two families, Wohlfarhtimonodaceae and Metaschinikowiceae, were more associated with the addition of streptomycin to media (Table 1), and not detected in the molecular controls (Figure 4). Streptomycin may have selected against the more abundant or more competitive organisms, allowing taxa from these families to grow.

Cultivation is critical to answer broad challenges of microbial ecology like deciphering microbial metabolisms or specific challenges like obtaining new isolates for antibiotic discovery. The study of pure cultures remains the best approach to comprehensively describe an organism’s physiological and metabolic properties, and efforts to optimize cultivation strategy have proven successful (13, 14, 16, 18, 21, 23, 28, 67–69). Our data adds to previous studies (26, 29) that also sought to assess how medium type and treatment could increase cultivated diversity, but adds the goal of “casting a wider net” to increase the diversity of bioactive isolates. We were able to add to our library of bioactive isolates and show that microbiomes from roadkill mammals are a useful source of bioactivity, as observed in Motley et al 2017. Lastly, we showed that through a thoughtful cultivation approach driven by bulk molecular analysis of cultivated taxa, we could cultivate a larger proportion of the diversity within an inoculum and even taxa not detected in the molecular controls. This work adds to the growing assessment of cultivation strategies using newer tools, like high-throughput sequencing, and shows how these methods can be applied to drug discovery efforts. Cultivation is influenced by many factors, and this work highlights some of those intricacies, like variability and stochasticity. Cultivation is complex and challenging but cultivating in combination with molecular tools can expand the observed diversity of an environment and its community.

## Acknowledgments

The authors thank Dr. H. Nunn and the team of Mammalian Microbiome undergraduate researchers (A. Byrd, J. Dewberry, B. Hansen, C. Hauger, J. Jacobs) for sample processing during cultivation. The authors also thank Dr. A. K. Dunn for helpful comments during the writing process. The isolates used for the pathogen panel were kindly provided by Dr. Robert Cichewicz, Institute for Natural Products and Research Technologies (INPART), University of Oklahoma.

## Supplementary Material

Figure S1: Bioassay plate showing three isolates generating a zone of inhibition within pathogen overlay.

Figure S2: Pie chart of 117 identified isolates at the genus level.

**Table S1.** A quasi-factorial approach using targeted media and treatments for gut microorganisms.

| Treatment/ Selection agents | Media |  |  |  |
| --- | --- | --- | --- | --- |
|  | 0.1 TSA | ROXY | 0.25 R2A | Blood Agar |
| None | X | X | X | X |
| Catalase | X | X | X |  |
| Hemin and alpha-ketoglutarate |  | X |  |  |
| 70% ethanol | X |  | X | X |
| Streptomycin | X |  | X | X |

Figure S3: Phylum-level characterization of molecular controls, triplicates shown for mouth (RC2M) and rectum (RC2R).

Figure S4: Bar chart describing the variability of consistent recovery of ASVs, data subset. Y axis indicates how many replicates of TSA plates in which the top 25 ASV were observed. Color corresponds to orifice.

Figure S5: Rarefaction curves showing plateaued sequencing depth of (a) cultivation samples and (b) molecular controls.

## References

1. Sogin ML, Morrison HG, Huber JA, Welch DM, Huse SM, Neal PR, Arrieta JM, Herndl GJ. 2006. Microbial diversity in the deep sea and the underexplored ‘‘rare biosphere’’. Proceedings of the National Academy of Sciences 103:12115–12120.

2. Donia MS, Cimermancic P, Schulze CJ, Brown LCW, Martin J, Mitreva M, Clardy J, Linington RG, Fischbach MA. 2014. A Systematic Analysis of Biosynthetic Gene Clusters in the Human Microbiome Reveals a Common Family of Antibiotics. Cell 158:1402–1414.

3. Tortorella E, Tedesco P, Palma Esposito F, January G, Fani R, Jaspars M, de Pascale D. 2018. Antibiotics from Deep-Sea Microorganisms: Current Discoveries and Perspectives. Marine Drugs 16:355–16.

4. Halfvarson J, Brislawn CJ, Lamendella R, Vázquez-Baeza Y, Walters WA, Bramer LM, D’Amato M, Bonfiglio F, McDonald D, González A, McClure EE, Dunklebarger MF, Knight R, Jansson JK. 2017. Dynamics of the human gut microbiome in inflammatory bowel disease. Nature Microbiology 2:1–7.

5. Arora-Williams K, Olesen SW, Scandella BP, Delwiche K, Spencer SJ, Myers EM, Abraham S, Sooklal A, Preheim SP. 2018. Dynamics of microbial populations mediating biogeochemical cycling in a freshwater lake. Microbiome 6:165–16.

6. Gilbert JA, Quinn RA, Debelius J, Xu ZZ, Morton J, Garg N, Jansson JK, Dorrestein PC, Knight R. 2016. Microbiome-wide association studies link dynamic microbial consortia to disease. Nature 535:94–103.

7. Singh BK, Bardgett RD, Smith P, Reay DS. 2010. Microorganisms and climate change: terrestrial feedbacks and mitigation options. Nature Reviews 8:779–790.

8. Staley JT, Konopka A. 1985. Meaurement of In Situ Activities of Nonphotosynthetic Microorganisms in Aquatic and Terrestrial Habitats. Annu Rev Microbiol 39:321–346.

9. Lloyd KG, Steen AD, Ladau J, Yin J, Crosby L. 2018. Phylogenetically Novel Uncultured Microbial Cells Dominate Earth Microbiomes. mSystems 3:431–12.

10. Eloe-Fadrosh EA, Ivanova NN, Woyke T, Kyrpides NC. 2016. Metagenomics uncovers gaps in amplicon-based detection of microbial diversity. Nature Microbiology 1:15032–4.

11. Schloss PD, Girard RA, Martin T, Edwards J, Thrash JC. 2016. Status of the Archaeal and Bacterial Census: an Update. mBio 7:995–10.

12. Nichols D, Cahoon N, Trakhtenberg EM, Pham L, Mehta A, Belanger A, Kanigan T, Lewis K, Epstein SS. 2010. Use of ichip for high-throughput in situ cultivation of “uncultivable” microbial species. Appl Environ Microbiol 76:2445–2450.

13. Stevenson BS, Eichorst SA, Wertz JT, Schmidt TM, Breznak JA. 2004. New strategies for cultivation and detection of previously uncultured microbes. Appl Environ Microbiol 70:4748–4755.

14. Connon SA, Giovannoni SJ. 2002. High-Throughput Methods for Culturing Microorganisms in Very-Low-Nutrient Media Yield Diverse New Marine Isolates. Appl Environ Microbiol 68:3878–3885.

15. Reasoner DJ, Geldreich EE. 1985. A New Medium for the Enumeration and Subculture of Bacteria from Potable Water. Appl Environ Microbiol 49:1–7.

16. Bartelme RP, Custer JM, Dupont CL, Espinoza JL, Torralba M, Khalili B, Carini P. 2020. Influence of Substrate Concentration on the Culturability of Heterotrophic Soil Microbes Isolated by High-Throughput Dilution-to-Extinction Cultivation. mSphere 5:489–15.

17. Davis KER, Joseph SJ, Janssen PH. 2005. Effects of Growth Medium, Inoculum Size, and Incubation Time on Culturability and Isolation of Soil Bacteria. Appl Environ Microbiol 71:826–834.

18. Henson MW, Pitre DM, Weckhorst JL, Lanclos VC, Webber AT, Thrash JC. 2016. Artificial Seawater Media Facilitate Cultivating Members of the Microbial Majority From the Gulf of Mexico. mSphere 1:e00028–16.

19. Kawasaki K, Kamagata Y. 2017. Phosphate-Catalyzed Hydrogen Peroxide Formation from Agar, Gellan, and κ-Carrageenan and Recovery of Microbial Cultivability via Catalase and Pyruvate. Appl Environ Microbiol 83:e01366–17.

20. Kim S, Kang I, Seo J-H, Cho J-C. 2019. Culturing the ubiquitous freshwater actinobacterial acI lineage by supplying a biochemical “helper” catalase. The ISME Journal 13:2252–2263.

21. Oberhardt MA, Zarecki R, Gronow S, Lang E, Klenk H-P, Gophna U, Ruppin E. 2015. Harnessing the landscape of microbial culture media to predict new organism-media pairings. Nature Communications 6:1–14.

22. Kopke B, Wilms R, Engelen B, Cypionka H, Sass H. 2005. Microbial Diversity in Coastal Subsurface Sediments: a Cultivation Approach Using Various Electron Acceptors and Substrate Gradients. Appl Environ Microbiol 71:7819–7830.

23. Browne HP, Forster SC, Anonye BO, Kumar N, Neville BA, Stares MD, Goulding D, Lawley TD. 2016. Culturing of “unculturable” human microbiota reveals novel taxa and extensive sporulation. Nature 533:543–546.

24. La Scola B, Khelaifia S, Lagier JC, Raoult D. 2014. Aerobic culture of anaerobic bacteria using antioxidants: a preliminary report. Eur J Clin Microbiol Infect Dis 33:1781–1783.

25. Dione N, Khelaifia S, La Scola B, Lagier JC, Raoult D. 2016. A quasi-universal medium to break the aerobic/anaerobic bacterial culture dichotomy in clinical microbiology. Clinical Microbiology and Infection 22:53–58.

26. Medina D, Walke JB, Gajewski Z, Becker MH, Swartwout MC, Belden LK. 2017. Culture Media and Individual Hosts Affect the Recovery of Culturable Bacterial Diversity from Amphibian Skin. Front Microbiol 8:32–15.

27. Justice NB, Sczesnak A, Hazen TC, Arkin AP. 2017. Environmental Selection, Dispersal, and Organism Interactions Shape Community Assembly in High-Throughput Enrichment Culturing. Appl Environ Microbiol 83:e01253-17-16.

28. Yang S-J, Kang I, Cho J-C. 2015. Expansion of Cultured Bacterial Diversity by Large-Scale Dilution-to-Extinction Culturing from a Single Seawater Sample. Microb Ecol 71:29–43.

29. Zehavi T, Probst M, Mizrahi I. 2018. Insights Into Culturomics of the Rumen Microbiome. Front Microbiol 9:D46–10.

30. Cummings LA, Hoogestraat DR, Rassoulian-Barrett SL, Rosenthal CA, Salipante SJ, Cookson BT, Hoffman NG. 2020. Comprehensive evaluation of complex polymicrobial specimens using next generation sequencing and standard microbiological culture. Sci Rep 10:1–12.

31. Hoffmann T, Krug D, Bozkurt N, Duddela S, Jansen R, Garcia R, Gerth K, Steinmetz H, Muller R. 2018. Correlating chemical diversity with taxonomic distance for discovery of natural products in myxobacteria. Nature Communications 9:1–10.

32. Helfrich EJN, Vogel CM, Ueoka R, Schafer M, Ryffel F, Muller DB, Probst S, Kreuzer M, Piel J, Vorholt JA. 2018. Bipartite interactions, antibiotic production and biosynthetic potential of the *Arabidopsis* leaf microbiome. Nature Microbiology 3:909–919.

33. Motley JL, Stamps BW, Mitchell CA, Thompson AT, Cross J, You J, Powell DR, Stevenson BS, Cichewicz RH. 2017. Opportunistic Sampling of Roadkill as an Entry Point to Accessing Natural Products Assembled by Bacteria Associated with Non-anthropoidal Mammalian Microbiomes. J Nat Prod 80:598–608.

34. van der Meij A, Worsley SF, Hutchings MI, van Wezel GP. 2017. Chemical ecology of antibiotic production by actinomycetes. FEMS Microbiology Reviews 41:392–416.

35. Stierle AA, Stierle DB, Kelley K. 2006. Berkelic Acid, A Novel Spiroketal with Selective Anticancer Activity from an Acid Mine Waste Fungal Extremophile. Journal of Organic Chemistry 71:5357–5360.

36. Ding Z-G, Li M-G, Zhao J-Y, Ren J, Huang R, Xie M-J, Cui X-L, Zhu H-J, Wen M-L. 2010. Naphthospironone A: An Unprecedented and Highly Functionalized Polycyclic Metabolite from an Alkaline Mine Waste Extremophile. Chem Eur J 16:3902–3905.

37. Rangseekaew P, Pathom-aree W. 2019. Cave Actinobacteria as Producers of Bioactive Metabolites. Front Microbiol 10:381–11.

38. Smanski MJ, Schlatter DC, Kinkel LL. 2015. Leveraging ecological theory to guide natural product discovery. Journal of Industrial Microbiology & Biotechnology 43:115–128.

39. Schatz A, Bugle E, Waksman SA. 1944. Streptomycin, a substance exhibiting antibiotic activity against Gram-positive and Gram-negative bacteria. Experimental Biology and Medicine 55:66–69.

40. Parada AE, Needham DM, Fuhrman JA. 2015. Every base matters: assessing small subunit rRNA primers for marine microbiomes with mock communities, time series and global field samples. Environ Microbiol 18:1403–1414.

41. Kraus EA, Beeler SR, Mors RA, Floyd JG, GeoBiology 2016, Stamps BW, Nunn HS, Stevenson BS, Johnson HA, Shapiro RS, Loyd SJ, Spear JR, Corsetti FA. 2018. Microscale Biosignatures and Abiotic Mineral Authigenesis in Little Hot Creek, California. Front Microbiol 9:544–13.

42. Caporaso JG, Kuczynski J, Stombaugh J, Bittinger K, Bushman FD, Costello EK, Fierer N, Peña AG, Goodrich JK, Gordon JI, Huttley GA, Kelley ST, Knights D, Koenig JE, Ley RE, Lozupone CA, McDonald D, Muegge BD, Pirrung M, Reeder J, Sevinsky JR, Turnbaugh PJ, Walters WA, Widmann J, Yatsunenko T, Zaneveld J, Knight R. 2010. QIIME allows analysis of high-throughput community sequencing data. Nat Methods 7:335–336.

43. Callahan BJ, McMurdie PJ, Rosen MJ, Han AW, Johnson AJA, Holmes SP. 2016. DADA2: High-resolution sample inference from Illumina amplicon data. Nat Methods 13:581–583.

44. Quast C, Pruesse E, Yilmaz P, Gerken J, Schweer T, Yarza P, Peplies J, Glöckner FO. 2013. The SILVA ribosomal RNA gene database project: improved data processing and web-based tools. Nucleic Acids Research 41:D590–6.

45. Yilmaz, P, Parfrey, LW, Yarza, P, Gerken, J, Pruesse, E, Quast, C, Schweer, T, Peplies, J, Ludwig, W, Glöckner, FO. 2014. Nucleic Acids Research 42:D643–8.

46. McMurdie PJ, Holmes S. 2013. phyloseq: an R package for reproducible interactive analysis and graphics of microbiome census data. PLoS ONE 8:e61217.

47. Dixon P. 2003. VEGAN, a package of R functions for community ecology. Journal of Vegetation Science 14:927–930.

48. De Cáceres M, Legendre P. 2009. Associations between species and groups of sites: indices and statistical inference. Ecology 90:3566–3574.

49. De Cáceres M, Sol D, Lapiedra O, Legendre P. 2011. A framework for estimating niche metrics using the resemblance between qualitative resources. Oikos 120:1341–1350.

50. Robinson JT, Thorvaldsdóttir H, Winckler W, Guttman M, Lander ES, Getz G, Mesirov JP. 2011. Integrative genomics viewer. Nature Biotechnology 29:24–26.

51. Kumar S, Stecher G, Li M, Knyaz C, Tamura K. 2018. MEGA X: Molecular Evolutionary Genetics Analysis across Computing Platforms. Molecular Biology and Evolution 35:1547–1549.

52. Newman DJ, Cragg GM. 2016. Natural Products as Sources of New Drugs from 1981 to 2014. J Nat Prod 79:629–661.

53. 2019. Antibiotic Resistance and Threats in the United States 2019. Center for Disease Control (CDC).

54. Clardy J, Fischbach MA, Walsh CT. 2006. New antibiotics from bacterial natural products. Nat Biotech 24:1541–1550.

55. Guo J, Fu X, Liao H, Hu Z, Long L, Yan W, Ding Y, Zha L, Guo Y, Yan J, Chang Y, Cai J. 2016. Potential use of bacterial community succession for estimating post-mortem interval as revealed by high-throughput sequencing. Sci Rep 6:1–11.

56. Metcalf JL, Wegener Parfrey L, González A, Lauber CL, Knights D, Ackermann G, Humphrey GC, Gebert MJ, Van Treuren W, Berg-Lyons D, Keepers K, Guo Y, Bullard J, Fierer N, Carter DO, Knight R. 2013. A microbial clock provides an accurate estimate of the postmortem interval in a mouse model system. eLife 2:403–19.

57. Jensen, V. 1968. The plate count technique. In TRG Gray & D Parkinson (Eds.), The ecology of soil bacteria (pp. 158–170). Liverpool University Press.

58. Hamdache A, Lamarti A, Aleu J, Collado IG. 2011. Non-peptide Metabolites from the Genus Bacillus. J Nat Prod 74:893–899.

59. Sumi CD, Yang BW, Yeo I-C, Hahm YT. 2015. Antimicrobial peptides of the genus Bacillus: a new era for antibiotics. Can J Microbiol 61:93–103.

60. Raaijmakers JM, Weller DM, Thomashow LS. 1997. Frequency of Antibiotic-Producing *Pseudomonas* spp. in Natural Environments. Appl Environ Microbiol 63:881–887.

61. Fuller AT, Mellows G, Woolford M, Banks GT, Barrow KD, Chain EB. 1971. Pseudomonic Acid: an Antibiotic produced by *Pseudomonas fluorescens*. Nature 234:416–417.

62. Hibbing ME, Fuqua C, Parsek MR, Peterson SB. 2009. Bacterial competition: surviving and thriving in the microbial jungle. Nature Reviews Microbiology 8:15–25.

63. de Vos MGJ, Zagorski M, McNally A, Bollenbach T. 2017. Interaction networks, ecological stability, and collective antibiotic tolerance in polymicrobial infections. Proceedings of the National Academy of Sciences 114:10666–10671.

64. Chao L, Levin BR. 1981. Structured habitats and the evolution of anticompetitor toxins in bacteria. Proceedings of the National Academy of Sciences 78:6324–6328.

65. Albright MBN, Chase AB, Martiny JBH. 2019. Experimental Evidence that Stochasticity Contributes to Bacterial Composition and Functioning in a Decomposer Community. mBio 10:247–13.

66. Traxler MF, Watrous JD, Alexandrov T, Dorrestein PC, Kolter R. 2013. Interspecies Interactions Stimulate Diversification of the *Streptomyces coelicolor* Secreted Metabolome. mBio 4:e00459–13.

67. Marmann A, Aly A, Lin W, Wang B, Proksch P. 2014. Co-Cultivation—A Powerful Emerging Tool for Enhancing the Chemical Diversity of Microorganisms. Marine Drugs 12:1043–1065.

68. Lagier J-C, Hugon P, Khelaifia S, Fournier P-E, La Scola B, Raoult D. 2015. The rebirth of culture in microbiology through the example of culturomics to study human gut microbiota. Clin Microbiol Rev 28:237–264.

69. Berdy B, Spoering AL, Ling LL, Epstein SS. 2017. In situ cultivation of previously uncultivable microorganisms using the ichip. Nat Protoc 12:2232–2242.

