## Supplementary figures and images for "Assessing and Maximizing Cultivated Diversity with Plate-Wash PCR and High Throughput Sequencing"

### Figure S1

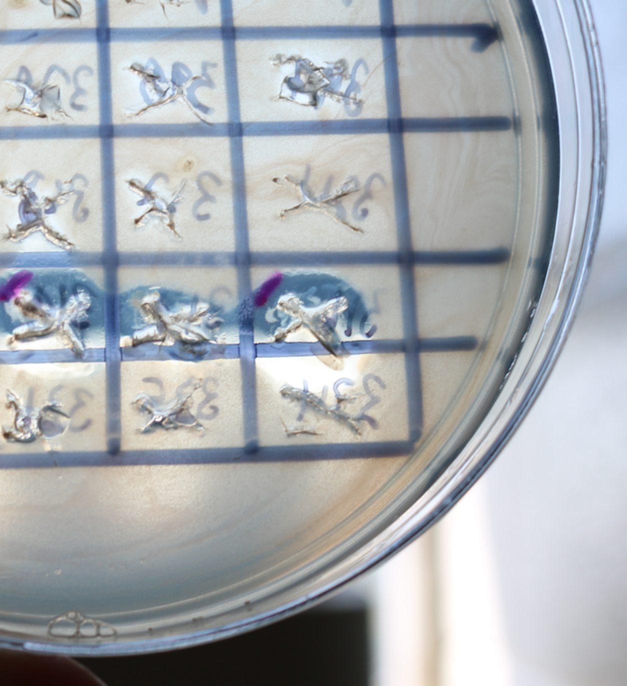

### Figure S2

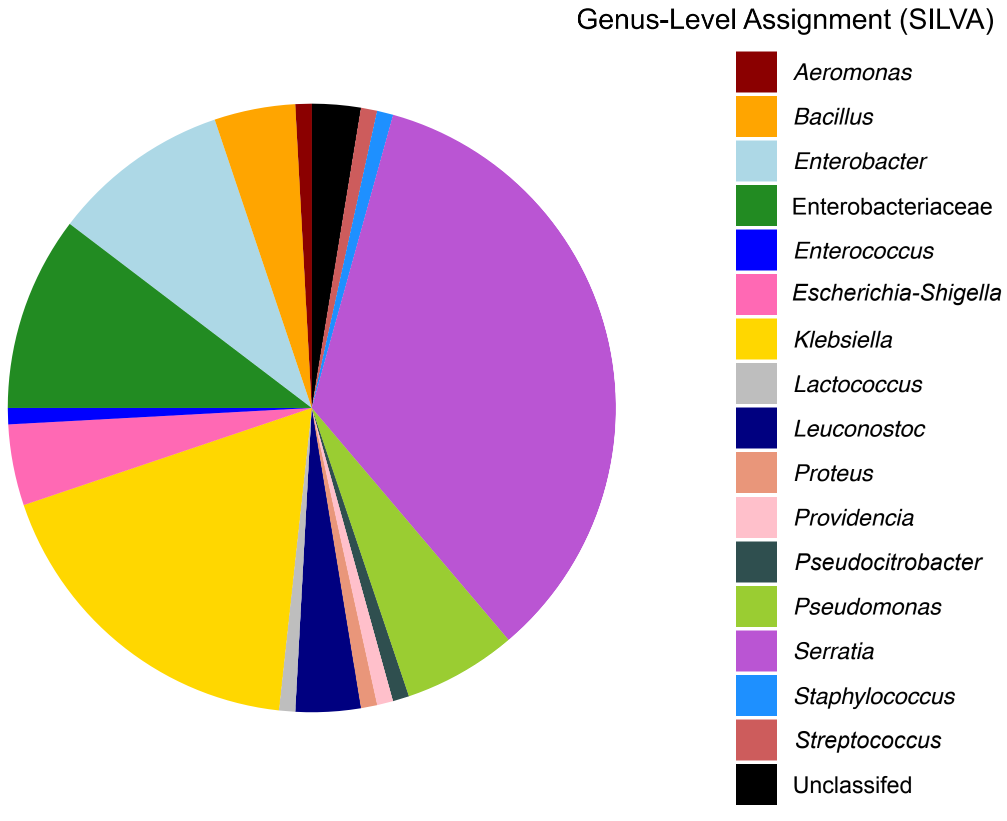

### Figure S3

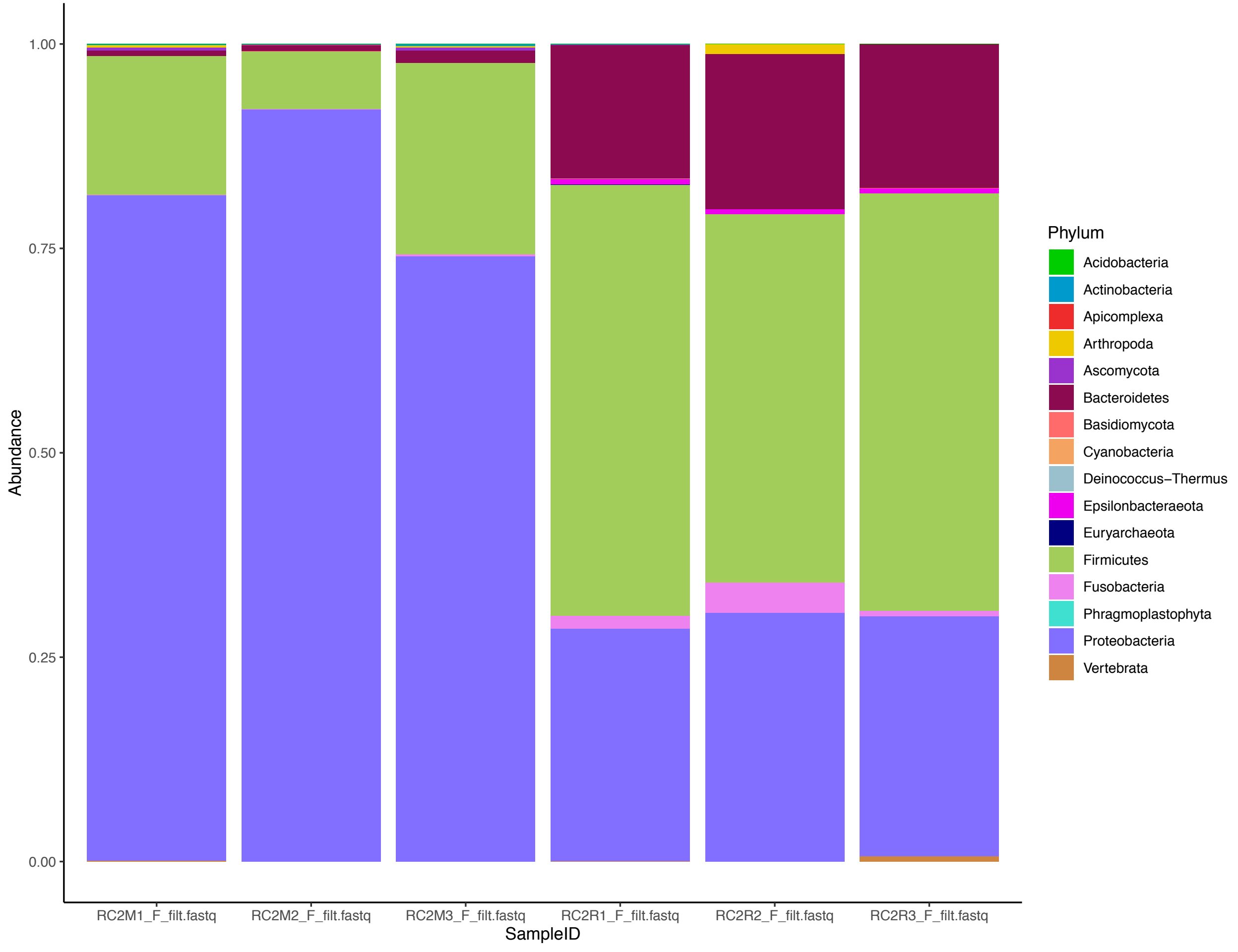

### Figure S4

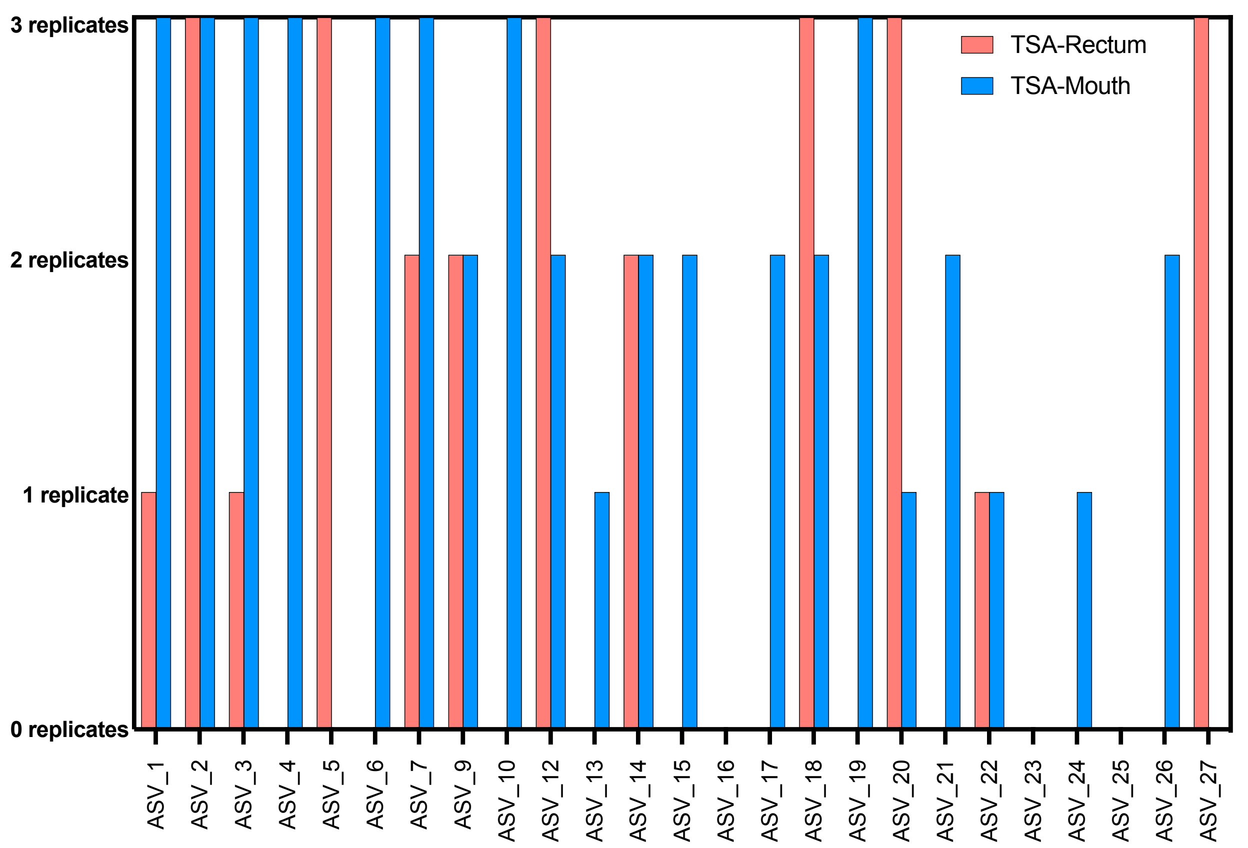

### Figure S5

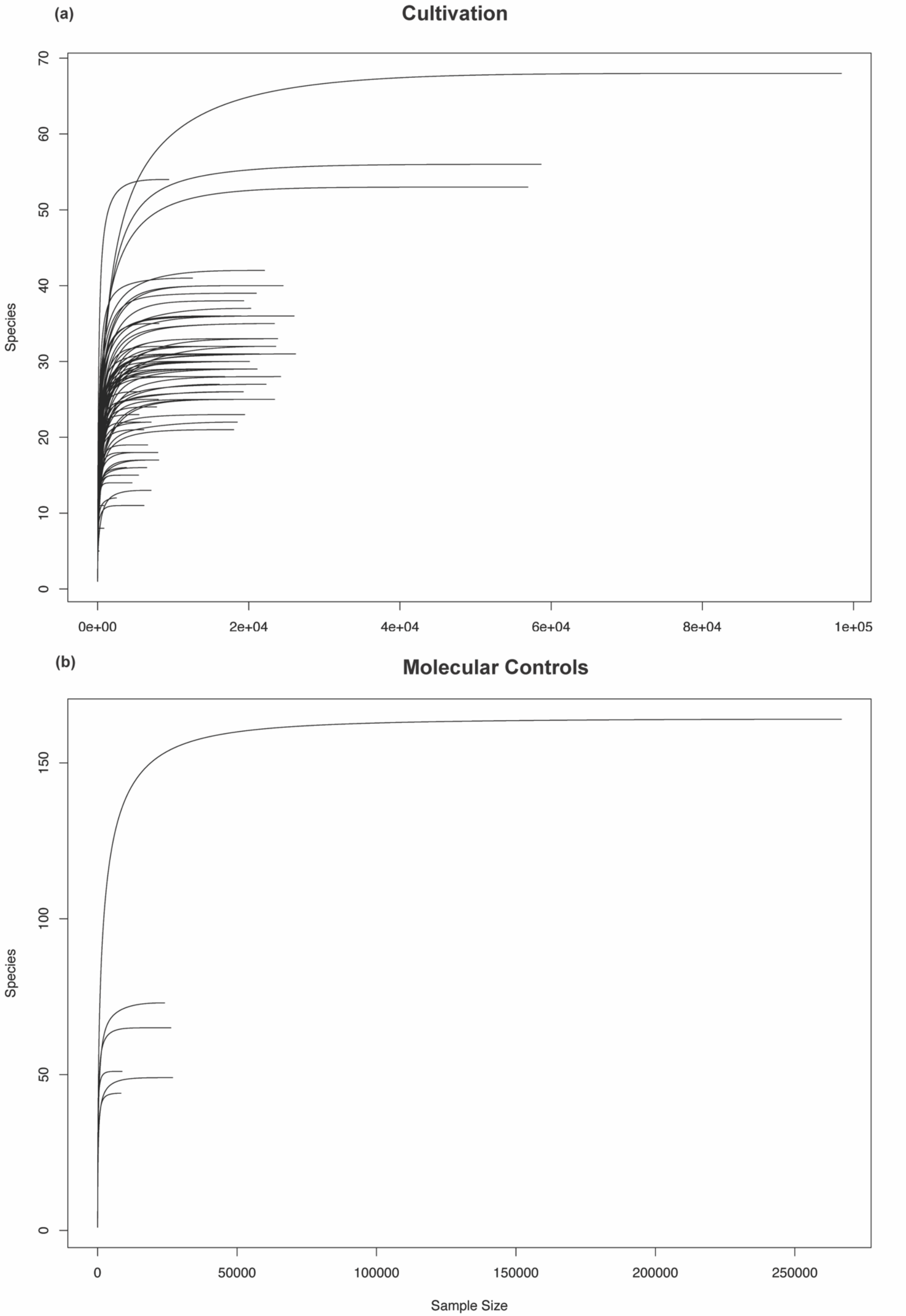
