## Supplementary material for "Assessing and Maximizing Cultivated Diversity with Plate-Wash PCR and High Throughput Sequencing": Table S1

| Table S1. A quasi-factorial approach using targeted media and treatments for gut microorganisms. | | | | |
| --- | --- | --- | --- | --- |
|  | **Media** | | | |
| Treatment/ Selection agents | 0.1 TSA | ROXY | 0.25 R2A | Blood Agar |
| None | X | X | X | X |
| Catalase | X | X | X |  |
| Hemin and alpha-ketoglutarate |  | X |  |  |
| 70% ethanol | X |  | X | X |
| Streptomycin | X |  | X | X |
